# Metabolic–epigenetic coupling between leucine catabolism and glycolysis drives CDK4/6 inhibitor resistance

**DOI:** 10.64898/2026.09.23.752387

**Authors:** Michael UJ Oliphant, Klarisa Norton, Kiran Kurmi, Howard Yang, Shakchhi Joshi, Jonah Lee, Nina Kozlova, Taru Muranen, John G Clohessy, Marcia Haigis, Senthil K Muthuswamy

## Abstract

Resistance to CDK4/6 inhibitors limits the durability of therapy for ER+ breast cancer. Despite the identification of mechanisms that regulate resistance, the metabolic adaptations that enable therapeutic escape remain poorly understood. Here, we identify a metabolic–epigenetic circuit that drives resistance by coordinately rewiring amino acid and glucose metabolism. CDK4/6 inhibitor–resistant ER+ tumor cells upregulate the leucine transporter SLC7A5, enhancing leucine uptake. SLC7A5 overexpression is sufficient to confer palbociclib resistance across ER+ cell lines, patient-derived organoids and xenografts. Stable isotope tracing in cell lines and in xenograft tumors revealed that leucine is catabolized through BCAT2 and HMGCL to increase acetyl-CoA levels, and elevated acetyl-CoA promotes H3K27 acetylation at the GLUT1 promoter, upregulating GLUT1 expression and glycolytic activity. Disrupting leucine transport, catabolism, or availability suppresses GLUT1 expression and restores therapeutic sensitivity in resistant models. In patients receiving palbociclib-based therapy, high SLC7A5 expression and coordinated SLC7A5–GLUT1 co-expression are associated with shorter progression-free survival. Together, these findings define a metabolic– epigenetic mechanism linking branched-chain amino acid catabolism to glycolysis and identify a biomarker-associated metabolic vulnerability in advanced ER+ breast cancer.

**Statement of Significance:** Resistance to CDK4/6 inhibitors is nearly universal in ER+ breast cancer. We identify a metabolic– epigenetic circuit in which acetyl-coA derived from leucine catabolism promotes epigenetic changes at the GLUT1 promoter, thereby increasing glucose uptake and driving glycolysis. Disrupting leucine transport, catabolism, or availability suppresses this program and restores drug sensitivity, identifying crosstalk between amino acid metabolism and glycolysis in regulating drug resistance. High expression of the leucine transporter, SLC7A5, is associated with shorter progression-free survival, revealing a biomarker-associated metabolic vulnerability.

## Introduction

Metabolic adaptation enables cancer cells to withstand microenvironmental stress while sustaining growth and proliferation^1–4^. Rewiring of metabolic pathways not only supports biosynthetic and bioenergetic demands but also shapes therapeutic response^5,6^. For example, alterations in glycolytic flux and redox homeostasis modulate sensitivity to first-line chemotherapies, underscoring a central role for metabolic state in determining drug response^6–8^. These observations have catalyzed efforts to therapeutically target tumor metabolism, with more than 20 metabolic inhibitors currently in clinical development or incorporated into combination regimens for advanced malignancies^5,9–12^. Despite this progress, how metabolic pathways are reprogrammed in cancer cells to regulate drug resistance remains incompletely understood.

Estrogen receptor–positive (ER+) breast cancer is the most prevalent breast cancer subtype^13^. Although therapies targeting ER signaling have substantially improved clinical outcomes, acquired resistance remains a major obstacle to durable disease control^14^. We previously demonstrated that the polarity protein LLGL2 promotes tamoxifen resistance in ER+ breast cancer by facilitating the cell-surface localization of the leucine transporter SLC7A5, thereby increasing leucine uptake and rewiring cellular metabolism^15^. High SLC7A5 expression has been associated with poor clinical outcomes across multiple solid tumor types, including breast, lung, gastric, and hepatocellular cancers, highlighting the clinical significance beyond ER+ breast cancer^16–18^. However, how increased leucine uptake rewires cellular metabolism to regulate drug resistance is unclear.

More recently, cyclin-dependent kinase 4 and 6 inhibitors (CDK4/6i) have become a standard-of-care treatment for endocrine-resistant breast cancer, significantly extending progression-free and overall survival^19–23^. Nevertheless, resistance to CDK4/6 inhibition invariably emerges, limiting long-term therapeutic benefit and underscoring the need to define the adaptive mechanisms that sustain tumor growth under CDK4/6 blockade^21–24^.

Here, we investigated whether SLC7A5 contributes to resistance to CDK4/6 inhibition in ER+ breast cancer. We find that SLC7A5 overexpression is sufficient to confer resistance to palbociclib in ER+ breast cancer cell lines, patient-derived organoids and xenografts. Mechanistically, elevated SLC7A5 levels enhance leucine uptake and catabolism, thereby increasing intracellular acetyl-CoA levels. Acetyl-CoA, in turn, promotes histone acetylation at the GLUT1 (SLC2A1) promoter, inducing GLUT1 expression, augmenting glucose uptake and glycolytic activity, and sustaining proliferation under CDK4/6 blockade. These findings define a metabolic–epigenetic axis in which leucine catabolism fuels chromatin remodeling to reprogram glucose metabolism, establishing a mechanistic cross-talk between branched-chain amino acid utilization and glycolysis in drug resistance.

## Results

### SLC7A5 promotes CDK4/6 inhibitor-resistant ER+ breast cancer

Cyclin-dependent kinase 4 and 6 inhibitors (CDK4/6i) have become a standard treatment for endocrine-resistant breast cancer, significantly increasing progression-free and overall survival^19–23^. However, resistance to CDK4/6 inhibition inevitably develops, limiting long-term treatment benefits and highlighting the need to identify mechanisms that sustain tumor growth under CDK4/6 blockade^21–24^. Analysis of transcriptomic data from tumors obtained from patients treated with CDK4/6 inhibitors^25^ show a significant increase in SLC7A5 expression in tumors post-treatment (which includes tumors that progressed) (Fig. 1a). This contrasted with four other amino acid transporters that we had previously reported to interact with LLGL2 in ER+ breast cancer cells^15^ (Fig. 1a). Furthermore, biomarker analyses from the PALOMA-2 and PALOMA-3 trials^26^ identified SLC7A5 among the genes most strongly associated with progression-free survival following CDK4/6 inhibitor therapy suggesting an enrichment of SLC7A5 regulated mechanisms during resistance in patients. Consistent with prior reports linking SLC7A5 to tamoxifen resistance and poor outcome in ER+ breast cancer^15,27^, we observed a marked increase in both SLC7A5 mRNA and cell-surface protein levels in an acquired palbociclib-resistant model (PalbR) (Fig. 1b,c) compared to sensitive counterparts (PalbS), suggesting selective enrichment of SLC7A5 during resistance evolution. We also found that PalbR cells exhibited higher intracellular branched-chain amino acid (BCAA) levels than PalbS cells (Fig. 1d), consistent with enhanced leucine metabolic flux.

**Figure 1.**
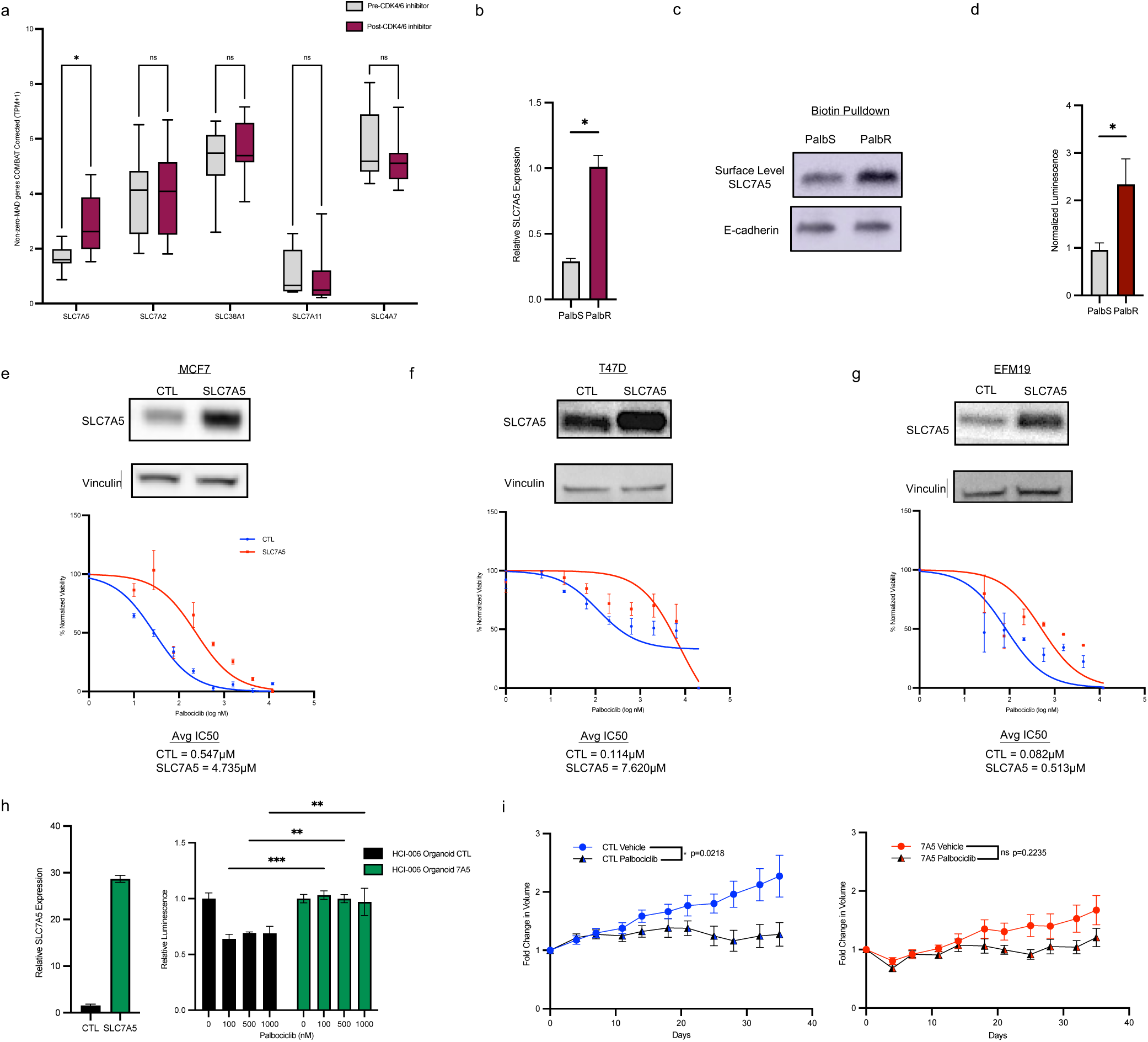
SLC7A5 promotes CDK4/6 inhibitor-resistant ER+ breast cancer. **a)** Expression of solute transporters in ER+ breast tumors from patients pre- and post-treatment with CDK4/6 inhibitors. **b)** mRNA and **c)** cell surface level expression of SLC7A5 in MCF7 palbociclib sensitive (PalbS) and acquired palbociclib resistant (PalbR) cells. **d)** Intracellular branched-chain amino acid levels in PalbS and PalbR cells. **e-g)** Control (CTL) and SLC7A5 overexpressing (7A5, or SLC7A5) MCF7, T47D, and EFM19 cells immunoblotted for SLC7A5 and Vinculin as a loading control. The graphs show the responses of all three cell lines treated with increasing doses of palbociclib, with the average (Avg) inhibitory concentration 50 (IC_50_) for each cell line indicated below the graphs. **h)** RT-qPCR of SLC7A5 mRNA expression and response to palbociclib treatment in HCI-006-tumor-derived control (HCI-006 Organoid CTL) and SLC7A5 overexpressing (HCI-006 Organoid 7A5) organoid lines. **i)** Tumor growth in mice transplanted with MCF7 control (CTL) or SLC7A5 overexpressing (SLC7A5) cells treated with vehicle or palbociclib (50 mg/kg). Data represent mean tumor volume ± SEM (n = 10 (CTL-Vehicle); 9 CTL-Palbociclib); 9 (7A5 Vehicle); and 7 (7A5 Palbociclib). Statistical significance was determined by two-way ANOVA at the endpoint (Day 35 post-treatment start).

To determine whether SLC7A5 is sufficient to confer resistance to CDK4/6 inhibition, we generated three independent ER+ cell models overexpressing SLC7A5 and exposed them to escalating doses of palbociclib. Across multiple ER+ cell lines, SLC7A5 overexpression (SLC7A5) conferred resistance to palbociclib, with an average ∼30-fold increase in IC_50_ (Fig. 1e-g) compared to control vector (CTL) transfected cells. Consistent with these findings, SLC7A5 overexpression (7A5) in patient-derived ER+ organoids significantly attenuated the response to palbociclib (Fig. 1h) compared to control vector-transfected (CTL) organoids, indicating that SLC7A5 is sufficient to promote resistance in clinically relevant tumor organoid models. To evaluate if SLC7A5 overexpression confers resistance *in vivo* MCF7 xenografts were generated from cells expressing either SLC7A5 or a control vector (CTL). Whereas control tumors were significantly growth-suppressed by palbociclib (p = 0.0218), SLC7A5-overexpressing tumors were refractory to treatment (p = 0.2235) (Fig. 1i). Importantly, baseline growth rates under vehicle conditions did not differ between CTL and SLC7A5-overexpressing tumors (p = 0.4184), indicating that SLC7A5 does not enhance intrinsic proliferative capacity but instead confers a selective advantage under CDK4/6 blockade. Together, these data establish that SLC7A5 overexpression is sufficient to drive robust and durable resistance to CDK4/6 inhibition across cell lines, organoids, and *in vivo* models.

### Leucine uptake is associated with activation of a glycolytic program

To determine whether amino acid metabolism is linked to broader metabolic reprogramming, we first assessed pathway-level changes associated with increased leucine uptake in SLC7A5-overexpressing cells (SLC7A5) (Fig 2a). Transcriptomic (RNA-seq) and metabolomic profiling of SLC7A5 and control cells (Fig 2a) revealed an unexpected enrichment of glycolysis as the top-ranked pathway (Fig. 2b and Extended Data Fig. 1a,b). Independent pathway analysis of SLC7A5-correlated genes in ER+ primary tumors similarly identified glycolysis as one of the most significantly enriched pathways (Fig. 2c), supporting the clinical relevance of this association. Consistent with these transcriptional changes, steady-state metabolomics demonstrated increased levels of glycolytic intermediates, including glucose-6-phosphate, fructose-6-phosphate, pyruvate, and lactate in SLC7A5 overexpressing cells both *in vitro* and xenograft tumors, without significant changes in tricarboxylic acid (TCA) cycle metabolites (Fig 2d and Extended Data Fig. 1c). Concordant with the increase in glycolysis, SLC7A5 overexpressing cells showed increased glucose uptake (Fig. 2e). This association was also observed in an acquired model of resistance (MCF7 PalbR), where glucose uptake was increased compared to sensitive parental cells (Fig. 2e). To determine whether this increase in glucose uptake reflected altered transporter expression, we examined GLUT family of glucose transporters. Interestingly, CDK4/6 inhibition has been reported to suppress GLUT1 expression through Rb–E2F–dependent reduction of c-Myc and HIF-1α^28^, suggesting that glucose transporter expression is itself a target of adaptation to CDK4/6 blockade. In contrast, GLUT1 protein levels were selectively increased in SLC7A5 cells, whereas GLUT4 remained unchanged (Fig. 2f). This extended to EFM19 and T47D cells, where increased GLUT1 expression increased in response to SLC7A5 overexpression (Fig 2g). Analysis of ER+ human tumor datasets further revealed a significant positive correlation between SLC7A5 and GLUT1 expression (Extended Data Fig. 1d), and their co-expression was associated with reduced overall survival in ER+ but not ER-negative disease (Fig 2h and Extended Fig 2). Together, these findings identify a link between leucine uptake, upregulation of GLUT1 expression, and activation of glycolysis.

**Figure 2.**
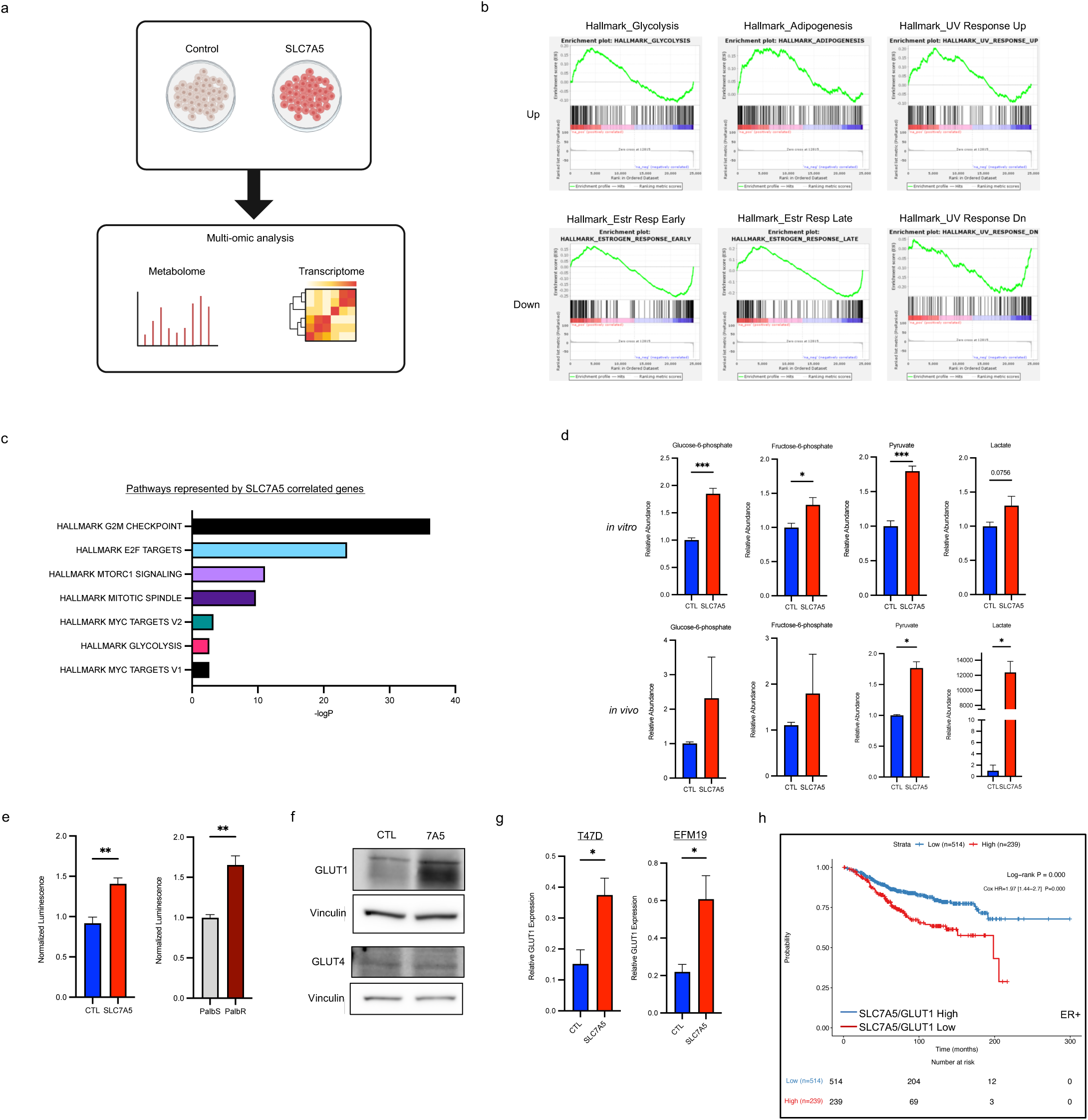
Leucine uptake is associated with activation of a glycolytic program. **a)** Schematic of multi-omic analyses in control (CTL) and SLC7A5 overexpressing (SLC7A5 or 7A5) MCF7 cells. **b)** GSEA enrichment plots for Top 3 upregulated (Up) and downregulated (Down) pathways in MCF7 SLC7A5 vs CTL. **c)** Pathways represented by the top 100 genes significantly correlated with SLC7A5 overexpression in ER+ tumors in the TCGA database as determined by Metascape pathway enrichment analysis. **d)** Levels of glycolytic metabolites in CTL and SLC7A5 MCF7 cells *in vitro* and in tumors grown *in vivo*. **e)** Glucose uptake in CTL and SLC7A5, palbociclib-sensitive (PalbS), and acquired palbociclib-resistant (PalbR) MCF7 cells. **f)** Immunoblot analysis for expression of GLUT1 and GLUT4 in control (CTL) and SLC7A5 MCF7 cells. Vinculin expression was used as a loading control. **g)** qRT-PCR analysis of GLUT1 in CTL and SLC7A5 overexpressing T47D and EFM19 cells. **h)** Overall survival of patients with ER+ tumors with high or low co-expression of SLC7A5 and GLUT1.

### Leucine catabolism drives glycolytic reprogramming through BCAT2

To investigate whether leucine utilization directly contributes to glycolytic activation, we traced ^13^C_6_-leucine flux in SLC7A5- and control cells *in vitro* and in xenograft tumors. Isotopic enrichment analysis revealed increased incorporation of ^13^C into intermediates of the leucine catabolic pathway and downstream metabolites, including acetyl-CoA and glutamate, indicating enhanced leucine catabolic flux in SLC7A5 cells (Fig. 3a,b). At the protein level, we observed increased expression of branched-chain amino acid transaminase 2 (BCAT2), but not BCAT1, enzymes that catalyze the first step of leucine catabolism^29–32^(Fig. 3c). Functionally, BCAT2 knockdown or treatment with the selective SLC7A5 inhibitor JPH203, reduced GLUT1 expression in both PalbR and SLC7A5 cells (Fig. 3d-i), establishing BCAT2-dependent leucine catabolism and SLC7A5 activity as key regulators of glycolytic reprogramming. Consistent with this relationship, analysis of ER+ tumor datasets demonstrated that co-expression of SLC7A5, BCAT2, and GLUT1 is associated with reduced overall survival in breast cancer patients (Fig. 3j), suggesting that this metabolic axis is clinically relevant. These findings position leucine catabolic flux upstream of glycolytic gene expression.

**Figure 3.**
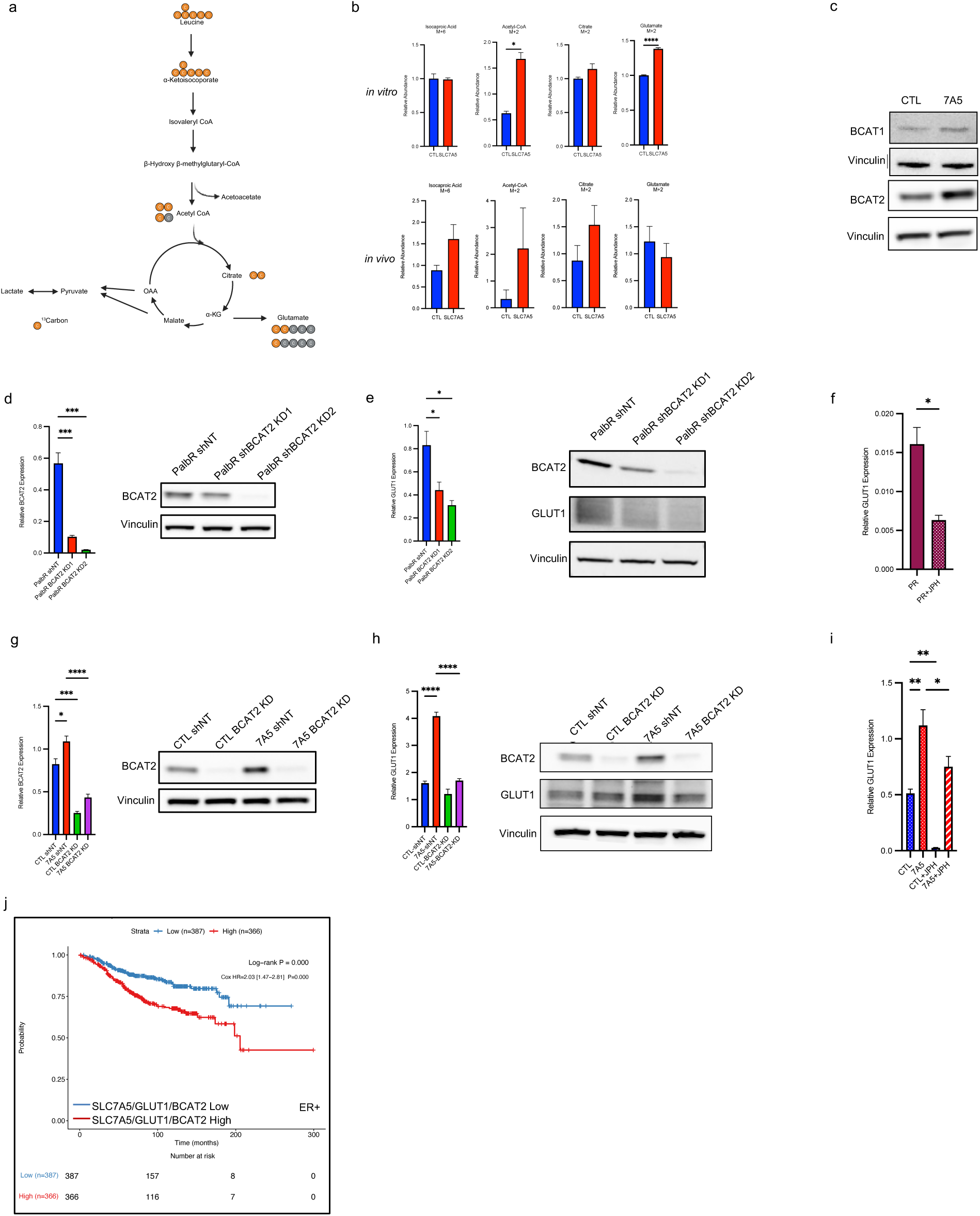
Leucine catabolism drives glycolytic gene expression through BCAT2. **a)** Schematic of the leucine catabolism pathway. **b)** Levels of ^13^C_6_ labelled leucine catabolism metabolites in control (CTL) and SLC7A5 overexpressing (SLC7A5 or 7A5) MCF7 cells *in vitro* and *in vivo*. **c)** Immunoblot analysis of BCAT1 and BCAT2 in CTL and 7A5 MCF7 cells. Vinculin expression was used as a loading control. **d**) qRT-PCR and immunoblot analysis of BCAT2 in palbociclib-resistant MCF7 cells expressing non-specific (PalbR-shNT) or two independent BCAT2 targeting (PalbR-BCAT2-KD1 and PalbR-BCAT2-KD2) short hairpin RNA (shRNA). **e)** qRT-PCR and immunoblot analysis of GLUT1 in palbociclib-resistant MCF7 cells expressing non-specific (PalbR-shNT) or two independent BCAT2 targeting (PalbR-BCAT2-KD1 and PalbR-BCAT2-KD2) short hairpin RNA (shRNA). **f)** qRT-PCR of GLUT1 in palbociclib-resistant MCF7 cells expressing treated with JPH203 (5μM) for 96h. **g)** qRT-PCR and immunoblot analysis of BCAT2 in CTL and 7A5 MCF7 cells with non-specific (CTL-shNT or 7A5-shNT) or BCAT2 KD2 targeting (CTL-BCAT2-KD; or 7A5-BCAT2-KD) short hairpin RNA (shRNA). **h)** qRT-PCR and immunoblot analysis of GLUT1 in CTL and 7A5 MCF7 cells with non-specific (CTL-shNT or 7A5-shNT) or BCAT2 KD2 targeting (CTL-BCAT2-KD; or 7A5-BCAT2-KD) short hairpin RNA (shRNA). **i)** qRT-PCR of GLUT1 in CTL and 7A5 MCF7 cells treated with JPH203 (5μM) for 96h. **j)** Overall survival in ER+ tumors with high or low co-expression of SLC7A5, GLUT1 and BCAT2.

### Leucine-derived acetyl-CoA links amino acid metabolism to chromatin regulation of glycolysis

To define the mechanism by which leucine catabolism regulates glycolytic gene expression, we examined downstream metabolites of leucine degradation. As shown above in leucine flux analysis, we observed a significant increase in ^13^C-labeled acetyl-CoA, a terminal product of leucine catabolism (Fig. 3). Because acetyl-CoA serves as the acetyl donor for histone acetylation, we hypothesized that increased acetyl-CoA may regulate transcription through chromatin modification. To test this, we engineered cells expressing a genetically encoded acetyl-CoA biosensor based on the bacterial PanZ protein fused to cytoplasmic or nuclear compartment-specific localization sequences and a circularly permuted GFP (cpGFP)^33^ (Fig. 4a). In this system, acetyl-CoA binding to PanZ promotes a conformational change and cpGFP fluorescence. SLC7A5 cells exhibited increased acetyl-CoA levels in both cytoplasmic and nuclear compartments, as indicated by an enhanced cpGFP signal (Fig. 4b). Interestingly, subcellular fractionation revealed the presence of BCAT2 and HMGCL in both nuclear and cytoplasmic compartments (Fig. 4c,d), suggesting a potential role for enzymes regulating acetyl-CoA generation from leucine catabolism in both compartments. Consistent with this, BCAT2 knockdown reduced acetyl-CoA levels in both compartments as measured by the biosensor (Fig. 4e) in SLC7A5 but not in control cells, demonstrating that leucine catabolism is required for the increase in nuclear and/or cytoplasmic acetyl-CoA observed in SLC7A5 cells. To determine whether leucine-derived acetyl-CoA is required for GLUT1 expression, we depleted HMGCL, the enzyme that catalyzes the final step of leucine degradation. HMGCL knockdown reduced GLUT1 mRNA expression (Fig. 4f), phenocopying BCAT2 depletion and identifying leucine-derived acetyl-CoA as a regulator of GLUT1 gene expression. SLC7A5 and PalbR cells exhibited increased global H3K27 acetylation (Fig. 4g), suggesting a relationship between leucine and gene expression. CUT&RUN analysis demonstrated an increase in H3K27ac occupancy at the GLUT1 promoter in SLC7A5 cells, which was reversed by BCAT2 depletion (Fig. 4h), indicating that leucine catabolism promotes histone acetylation to drive GLUT1 transcription. Together, these findings support a mechanism in which leucine-derived acetyl-CoA directly links amino acid metabolism to chromatin-mediated activation of glycolysis.

**Figure 4.**
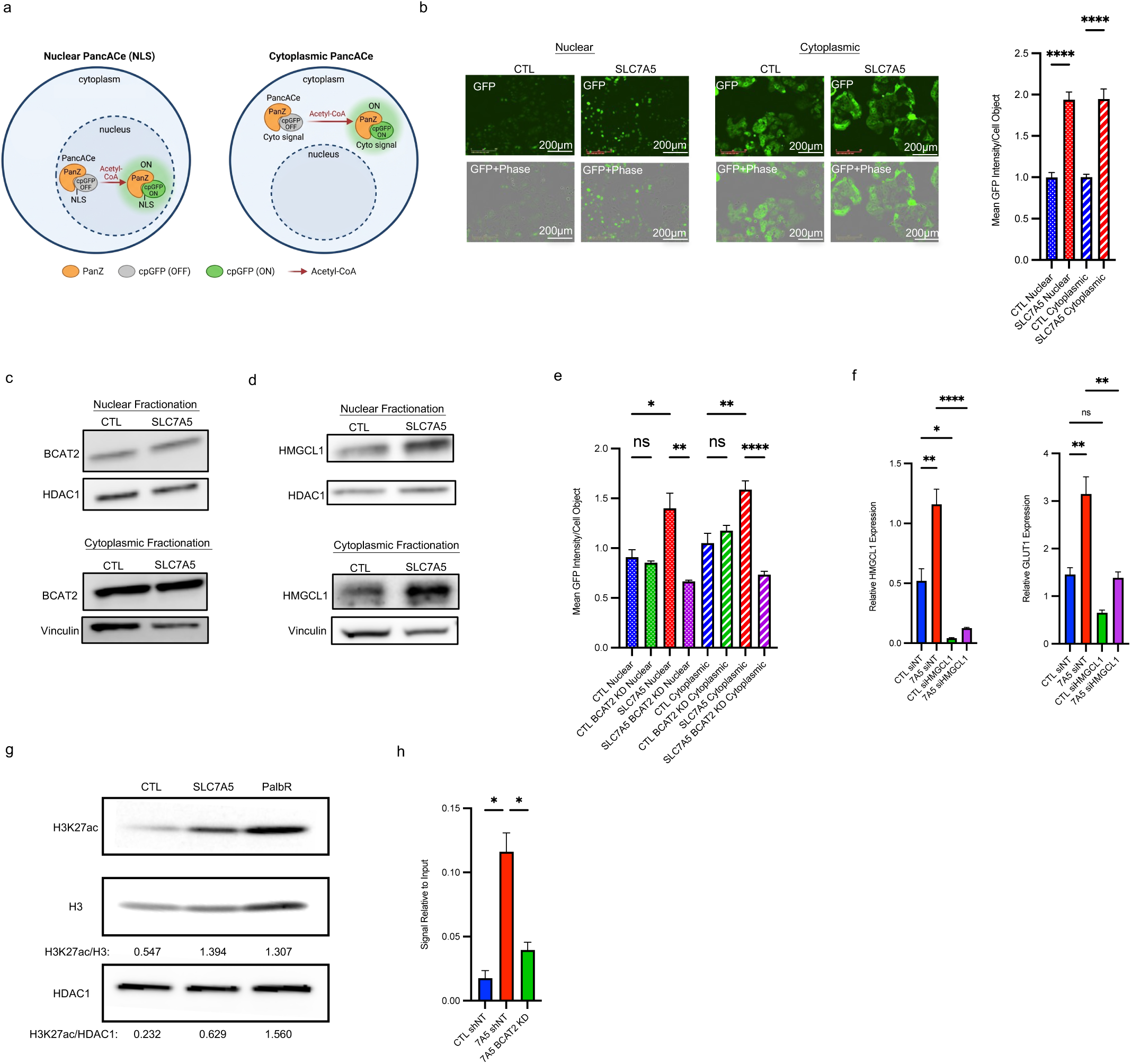
Leucine-derived acetyl-CoA epigenetically activates glycolytic transcription. **a)** Schematic of nuclear vs cytoplasmic localization of the acetyl-CoA biosensor. **b)** Images of nuclear and cytoplasmic GFP (acetyl-CoA) in control (CTL) and SLC7A5 overexpressing (SLC7A5 or 7A5) MCF7 cells (left) and GFP levels (right). **c)** Immunoblot analysis of BCAT2 in nuclear and cytoplasmic fraction from CTL and SLC7A5 MCF7 cells. **d)** Immunoblot analysis of HMGCL in nuclear and cytoplasmic fraction from CTL and SLC7A5 MCF7 cells. **e)** GFP (acetyl-CoA) levels in CTL and 7A5 MCF7 cells, with non-specific (CTL-shNT or 7A5-shNT) BCAT2 targeting (CTL-BCAT2-KD; or 7A5-BCAT2-KD) short hairpin RNA (shRNA). **f)** qRT-PCR analysis of HMGCL and GLUT1 in control and SLC7A5 overexpressing MCF7 cells with and without HMGCL KD. **g)** Immunoblot analysis of H3K27ac in CTL and SLC7A5 and acquired palbociclib resistant (PalbR) MCF7 cells. Quantification normalized to H3 and HDAC1 is shown below. **h)** CUT&RUN qPCR of GLUT1 promoter in CTL and 7A5 MCF7 cells, with CTL-shNT or 7A5-shNT or 7A5-BCAT2-KD shRNA.

### Leucine uptake and catabolism sustain proliferation under CDK4/6 inhibition

To determine whether leucine catabolism is required for proliferation under CDK4/6 inhibition, we tested the impact of disrupting leucine availability, transport, and catabolism in resistant ER+ breast cancer cells. Leucine restriction reduced the palbociclib IC_50_ in PalbR cells by ∼2.5-fold (Fig. 5a), linking leucine availability to resistance. Co-treatment of PalbR cells with palbociclib and the selective SLC7A5 inhibitor JPH203 produced an approximately threefold reduction in IC_50_ compared to either agent alone (Fig. 5b), and a functional synergy between CDK4/6 blockade and inhibition of leucine transport as determined by four synergy scoring models (Extended Data Fig. 3a). This resistance was dependent on leucine catabolism, as BCAT2 depletion restored drug sensitivity in both SLC7A5 overexpressing and PalbR cells (Fig. 5c,d).

**Figure 5.**
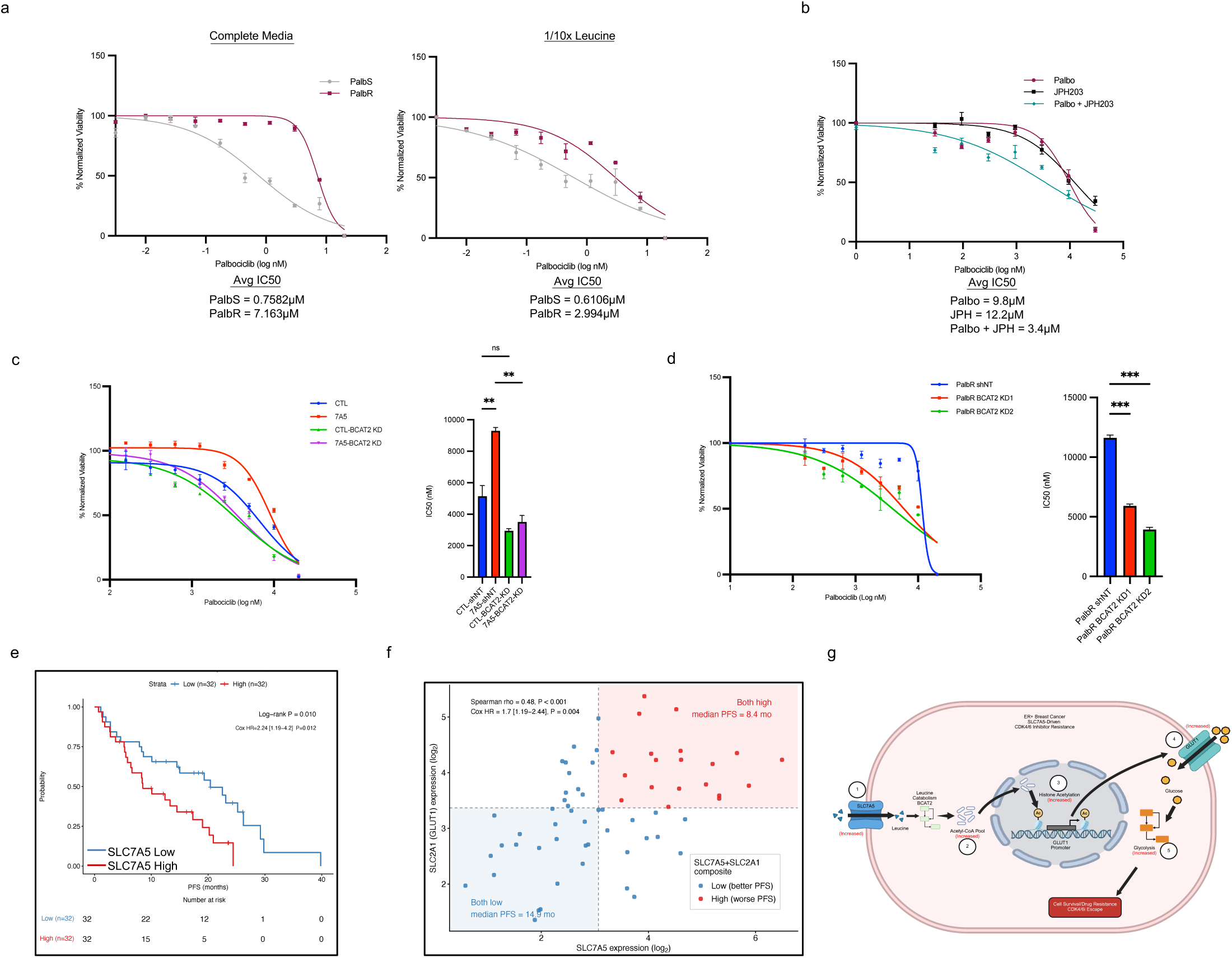
Leucine uptake and catabolism sustain proliferation under CDK4/6 inhibition. **a)** Palbociclib sensitivity for PalbS or PalbR MCF7 cells grown in media containing normal leucine concentration (1x Leucine) or in media containing 1/10^th^ the amount of leucine (1/10x Leucine). **b)** MCF7 PalbR cells treated with palbociclib, JPH203 (6.25µM), or palbociclib + JPH203. **c)** Palbociclib treatment dose curve (left), and IC_50_ values (right) in CTL and 7A5 MCF7 cells with non-specific (CTL-shNT or 7A5-shNT) or BCAT2 KD2 targeting (CTL-BCAT2-KD; or 7A5-BCAT2-KD) short hairpin RNA (shRNA). **d)** Palbociclib treatment dose curve (left), and IC_50_ values (right) in PalbR MCF7 cells expressing PalbR-shNT or PalbR-BCAT2-KD1 and PalbR-BCAT2-KD2 shRNA. **e)** Kaplan-Meier survival analysis of progression-free survival (PFS) stratified by SLC7A5 expression in patients with HR+/HER2− metastatic breast cancer receiving palbociclib (GSE186901; n=64). **f)** Scatterplot of baseline tumor RNA-seq expression (log₂ TPM) for SLC7A5 and SLC2A1 in the GSE186901 palbociclib cohort (n=64). Dashed lines indicate median expression thresholds for each gene. Points are colored by SLC7A5+SLC2A1 group. Patients with both genes above median (red) versus all others (blue). Spearman ρ=0.48, P<0.001. **g)** Model of SLC7A5-mediated resistance to CDK4/6 inhibition in ER+ breast cancer.

To assess whether our findings translate to patient outcomes, we analyzed a publicly available cohort of HR+/HER2− metastatic breast cancer patients treated with palbociclib plus endocrine therapy (GSE186901; n=64 baseline tumor biopsies)^34^. Patients with high SLC7A5 expression exhibited significantly shorter progression-free survival (PFS) compared to those with low expression (HR=2.24 [95% CI: 1.19–4.20], log-rank P=0.010; Fig. 5e). We next examined whether co-expression of SLC7A5 with GLUT1 defined a clinically distinct subgroup. SLC7A5 and GLUT1 expression were significantly correlated in palbociclib-treated tumors (Spearman ρ=0.48, P<0.001; Fig. 5f), and the composite high-expression group exhibited significantly decreased PFS (Cox HR=1.70 [1.17–2.19], P=0.004; median PFS 8.4 vs 14.9 months; Fig. 5f). Together, these preclinical and patient data establish that the leucine-driven metabolic-epigenetic program sustains proliferation under CDK4/6 blockade and identifies a coordinated metabolic signature associated with poor patient outcomes (Fig. 5g).

## Discussion

Resistance to targeted therapies has largely been interpreted within the framework of genetic adaptation and reactivation of signaling pathways. Our findings extend this view by identifying a mechanism by which metabolic pathways are directly coordinated at the chromatin level. Specifically, we show that leucine catabolism generates acetyl-CoA that promotes H3K27 acetylation at the GLUT1 promoter, thereby linking amino acid metabolism to transcriptional control of glycolysis. This mechanism expands current models of leucine biology, which have primarily focused on mTORC1 activation^35,36^. Although we have shown that leucine-dependent mTOR signaling is preserved in SLC7A5-overexpressing cells^15^, sensitivity to mTOR inhibition was unchanged in SLC7A5-overexpressing cells (Extended Data Fig. 3b) and SESTRIN2 expression was unaltered (Extended Data Fig. 3c), a determinant of mTOR inhibitor sensitivity^37^. Thus, our results show that an increase in intracellular leucine can serve as an independent transcriptionally instructive input in a catabolism-dependent manner. These findings are consistent with emerging evidence that metabolites can regulate chromatin state^38,39^, but extend this concept by demonstrating cross-regulation between distinct metabolic pathways.

The presence of BCAT2 and HMGCL in nuclear compartments suggests that acetyl-CoA production may be spatially coupled to transcriptional regulation, thereby integrating nutrient availability with gene expression. In this context, glycolytic activation is not simply a downstream consequence of oncogenic signaling, but can be imposed by upstream amino acid flux. While this metabolic crosstalk supports proliferation under CDK4/6 inhibition, the underlying mechanism is not specific to this therapeutic context. Instead, it reveals a broader principle whereby nutrient availability and metabolism can directly regulate transcriptional programs that govern other metabolic pathways. This may help explain the frequent occurrence of upregulation of amino acid transporters^40–43^ and glycolytic activation across cancers^4,44^. From a therapeutic perspective, modulating intracellular leucine levels, targeting leucine transport, or catabolism represents a strategy to disrupt this cross-regulatory axis. The observed synergy between CDK4/6 inhibition and SLC7A5 blockade supports this approach, although the broader implication is that metabolic dependencies may be best addressed by targeting upstream regulatory nodes rather than individual pathways.

## Methods

### Cell lines

MCF7, T47D, and EFM19 cells were cultured in Eagle’s minimum essential medium (EMEM) (ATCC 30-2003) supplemented with 10% FBS, 10 µg/ml insulin, and 1% penicillin/streptomycin (P/S) (Passaging Media). Palbociclib-resistant and sensitive MCF7 cells were kindly provided by Nicholas Turner (Institute of Cancer Research and Royal Marsden Hospital, London, UK.) and were cultured as previously described^45^. HCI-006 was grown as previously described in Breast Tumor Organoid Media (BTOM)^46^. All cell lines and organoids were monitored for mycoplasma by PCR and maintained in mycoplasma-free conditions.

### Dose response curves

On day zero, 3,000 cells per well were plated in a 96-well plate in passaging media. The media was changed to complete assay media (EMEM, 2% dialyzed FBS, and 1% P/S) 24 hours later. For 1/10th leucine media, leucine-free EMEM was prepared from EMEM without amino acids (M3859-01) by adding back all amino acids at standard EMEM concentrations, except L-leucine was added at 1/10th the standard concentration (0.04 mM). Cells were treated with indicated drugs (Palbociclib, Cat# S1579; JPH203, Cat# S8667; Everolimus, Cat# S1120; Abemaciclib, Cat# S5716; Ribociclib, Cat# S5188; Fulvestrant, Cat# S1191; all from Selleckchem) on day 1 and refreshed every other day for 5 days. Viability was assessed using the CyQuant cell proliferation assay (Thermo Fisher, Cat# C7026) according to the manufacturer’s protocol. All results were normalized to DMSO vehicle controls.

### Lentiviral vectors

The vector pLJM1-SLC7A5 was used to generate cells with SLC7A5 overexpression^47^. BCAT2 knockdown was achieved using pLKO.005 vectors (shGFP for control, BCAT2-KD1, and BCAT2_KD2). TRCN0000231747 was used for shGFP, TRCN0000286203 for BCAT2-KD1, and TRCN0000286266 for BCAT2_KD2. For acetyl-CoA reporters, Nuclear PancACe (Addgene 215706) and Cytoplasmic PancACe (Addgene 215708) were used. Each lentivirus was produced by transfection of HEK293 T cells with pCMV delta R8.2 (Addgene 12263) and pCMV-VSV-G (Addgene 8454) using polyethyleneimine (PEI, ThermoFisher 043896.01).

### Animal Studies

All animal experiments were performed in conjunction with the Preclinical Murine Pharmacogenetics Core (RRID: SCR_009675) at the Beth Israel Deaconess Medical Center and were approved by and in accordance with the guidelines of the Beth Israel Deaconess Medical Center Institutional Animal Care and Use Committee (animal protocol #077-2021-24). 90-day-release 17ß-estradiol pellets (0.5 mg; Innovative Research of America #NE-121) were implanted subcutaneously into six-week-old female Crl:NU(NCr)-Foxn1nu/nu mice (Charles River #490). Two days later, 5 x 10^6 15 MCF7 cells (CTL or SLC7A5) were suspended in 100µL of 1:1 PBS to Matrigel (Corning #356230) and injected into the left mammary fat pad of all mice. Tumors were measured twice per week with calipers, and volume was calculated with the formula (Width^2 * Length) / 2. Mice were enrolled in treatment when their average tumor volume reached 250 cubic mm. Mice were assigned to one of two treatment groups: palbociclib HCl (50 mg/kg; 0.5% methyl cellulose, 0.2% Tween-80, 99.3% normal saline; MedChemExpress #HY-50767C) daily by oral gavage, or vehicle daily by oral gavage. Mice bearing control tumors were treated for 28 days. Seven days after the last treatment, control tumor mice were euthanized by carbon dioxide asphyxiation, and tumor, lung, and liver tissue were harvested. One portion of each tissue was flash frozen, and another portion was fixed in 10% neutral-buffered formalin for 24 hours. Blood plasma was also collected from a subset of mice via post-mortem cardiac puncture using lithium heparin as the anticoagulant. OE tumor-bearing mice were treated for 35 days. Four days after the last treatment, OE tumor mice were euthanized and processed as described^47^. (N.B., the SLC7A5 tumor-bearing, vehicle-treated mice received both palbociclib vehicle as described and 5% DMSO/95% corn oil once daily by intraperitoneal injection from treatment day one through day 24)^48^.

### Live cell imaging

For real-time quantitative live-cell proliferation analysis, cells were seeded onto 96-well plates as outlined above in passage media. The next day, passage media were removed, the cells were washed 2x with PBS, replaced with assay media, and treated with drugs, if applicable. The plate was then moved into the IncuCyte® S3 System (Essen BioScience) for live phase contrast recording of cell confluence and GFP fluorescence (for Acetyl-CoA). The fluorescence and confluence analyses were performed using the basic IncuCyte software settings.

### Immunoblot

Immunoblotting was conducted as previously described^47^. Briefly, cells were lysed with lysis buffer (50 mM Tris-HCl, pH 7.5, 150 mM NaCl, 5 mM EDTA, 1% Triton X-100, protease inhibitors, and phosphoSTOP tablets (Roche)). The supernatant was subjected to SDS–PAGE, and proteins were transferred onto a PVDF membrane (Millipore #IPVH00010). After blocking with 1% BSA/TBS-0.1% Tween 20, the primary antibody was applied overnight at 4 °C. HRP-conjugated secondary antibodies (GE Healthcare) and HRP substrate (Pierce) were used for detection. Membranes were exposed to ImageQuant800 (GE Healthcare). Nuclear fractionation was done using the NE-PER™ Nuclear and Cytoplasmic Extraction kit (78833).

For assessment of SLC7A5 surface levels, cells were rinsed with ice-cold PBS and surface proteins were labelled with 400 µM EZ-link-sulfo-NHS-SS-Biotin/PBS for 30-60 min at 4 °C with gentle rocking. The reaction was quenched with 2 ml 150 mM glycine. After two rinses with ice-cold PBS, cells were lysed in lysis buffer. Fifty micrograms of protein lysate was incubated with 20 µl Streptavidin Sepharose High Performance Beads (GE Healthcare) and rotated for 1.5h at 4 °C. Streptavidin beads were washed three times with lysis buffer, and proteins were eluted with 2x sample buffer.

### Antibodies

Anti-E-CADHERIN (clone 24E10, Cat# 3195), anti-VINCULIN (Cat# 4650), anti-GLUT1 (Cat# 73015), anti-GLUT4 (Cat# 9001), anti-BCAT1 (clone D4V2T, Cat# 16431), anti-BCAT2 (clone D8K3O, Cat# 79764), anti-HMGCL (clone E5H7V, Cat# 39913), anti-H3K27ac (clone D5E4, Cat# 8173), anti-H3 (Cat# 9715), anti-HDAC1 (Cat# 2062), and anti-SESTRIN2 (clone D1B6, Cat# 8487) antibodies were purchased from Cell Signaling Technology. Anti-SLC7A5 (BMP011) antibody was purchased from MBL. Antibodies were used with 1/1,000 dilution for immunoblot.

### Organoid culture and SLC7A5 overexpression

Tumor chunks from the previously reported HCI-006 xenograft model^49^ were kindly provided by Alana Welm (Huntsman Cancer Institute, University of Utah). Organoids were established from tumor chunks, cultured, and treated with drugs as previously described^46^. SLC7A5 overexpression was achieved by infecting HCI-006 organoids with SLC7A5-expressing lentivirus. For lentiviral infection of organoids, a modified spinfection was performed. Pelleted organoids were resuspended in lentivirus media plus polybrene and centrifuged for 1h at 1000rpm, 37 °C. After centrifugation, organoids were plated as described above. Drug selection by puromycin (1ug/ml) was performed 72-96 hours post-infection for 3 days to establish stable overexpression.

### Survival Analysis and expression data

For survival data analyzed through Kaplan–Meier Plotter (http://kmplot.com/analysis/), we used the 201195_s_at probe for SLC7A5, 201249_at for GLUT1, 203576_at for BCAT2, 225285_at for BCAT1 (mean expression) to analyze the relationship between SLC7A5/GLUT1, SLC7A5/BCAT1, and SLC7A5/BCAT2 gene expression and overall survival. After removal of biased array data (patient and clinical heterogeneity, different outcome measures, and size effects) and redundant samples, results were split by the “auto select best cut off” and graphed to show the approximate low versus high expression and stated P values. For assessment of transporter expression from ER+ breast cancer patients pre- and post-CDK4/6i, we extracted and plotted the non-zero-MAD genes COMBAT corrected (TPM+1) values for each gene from Ref 25.

### Metabolomics analysis

For in vitro analysis, cells were grown to 70–80% confluence in complete assay media as outlined above for 72h in 6-well plates. Plates were washed twice with ice-cold DPBS, and metabolism was quenched by floating the plate in liquid nitrogen for 10 seconds. Plates were kept on dry ice and in a wet ice bath for all remaining steps. One milliliter of extraction solvent (80 percent LC-MS-grade methanol and 20 percent water with glutaric acid at 1 microgram per milliliter) was added per well, and the extract was transferred to Eppendorf tubes on dry ice until further processing. A mock extraction from a well containing only media was processed in parallel. Extracts were vortexed for 10 minutes in the cold room, centrifuged for 15 minutes at 13,300 rpm at 4°C, and the supernatant was transferred to new tubes without disturbing the protein pellet. Protein pellets were stored at −80°C for later protein quantification if necessary. Supernatants were dried in a refrigerated SpeedVac at 4°C and stored at −80°C until LC-MS analysis. Before LC-MS, dried extracts were reconstituted in a resuspension solvent (50 percent acetonitrile and 50 percent LC-MS water) containing D8-phenylalanine at 0.2 micrograms per milliliter. We targeted 5,000–10,000 cells per microliter at resuspension by adjusting seeding density and extraction volume. Parallel plates provided cell counts and total protein for normalization; results are reported normalized to cell number or protein, as indicated. For in vivo profiling, 1 million MCF7 control or SLC7A5 cells were injected into the mammary fat pad of NSG mice, and tumors were allowed to grow to an area of ∼150 mm^2^ before being excised and harvested. Tumors were homogenized with pre-cooled 80% methanol in HPLC-grade water (1 ml 80% methanol per 100 mg sample) in dry ice and further processed as outlined above. For leucine tracing experiments, in vitro cells were traced with uniformly labeled 13C6 Leucine (0.4 mM, CLM-2262-H, Cambridge Isotope Laboratories, Inc.) for 24 h in complete assay media, and in vivo (15mg/ml) for 15 min via 200 µl IP and mammary fat pad injection prior to tumor harvesting.

For steady-state and tracing studies, metabolites were separated using a Vanquish U-HPLC system coupled to a Q Exactive HF-X hybrid quadrupole-Orbitrap mass spectrometer (Thermo Fisher Scientific) equipped with a heated electrospray ionization (HESI) source operating in negative ion mode. Chromatographic separation was performed on an iHILIC column (5 µm, 150 × 2.1 mm; HILICON). The mobile phases consisted of solvent A (water containing 20 mM ammonium carbonate and 0.1% ammonium hydroxide) and solvent B (acetonitrile). The LC gradient was run at a flow rate of 0.15 mL/min as follows: 0–23 min, linear gradient from 95% B to 5% B; 23–25 min, hold at 5% B; 25– 25.5 min, gradient to 95% B at 0.20 mL/min; 25.5–32.5 min, hold at 95% B; and 32.5–33 min, return to 0.15 mL/min at 95% B for re-equilibration. Targeted feature extraction and metabolite quantification were performed using TraceFinder v4.1 (Thermo Fisher Scientific). Metabolite identification and peak area integration were based on accurate mass and retention time matched to an in-house library of authenticated standards. Data were normalized to cell number or total protein, and an internal standard was used to account for variability introduced during sample handling, preparation, and injection. Downstream statistical and pathway analyses were performed using the MetaboAnalyst 5.0 platform.

### RNAseq Analyses

The RNAseq reads were aligned to the human reference genome (hg38) with STAR. Gene expression quantification was performed using RSEM. Differentially expressed genes between MCF7 control and SLC7A5 cells were identified using DESeq2 from RSEM count data. For the pathway analysis, we ranked the entire gene list based on DESeq2 statistics (p-values and fold changes) and applied the Gene Set Enrichment Analysis (GSEA) software *GSEAPreranked* to compute Normalized Enrichment Scores (NES) and identify enriched Hallmark pathways.

### Glucose uptake and BCAA intracellular levels

Cells were first grown in complete assay medium as outlined above for 72h. For Glucose uptake, the medium was removed, washed with PBS 2X, and replaced with fresh complete assay medium containing 1 mM 2-DG. After 30 min incubation, glucose uptake was measured using the Glucose Uptake-Glo assay kit (Promega). For BCAA levels, cells were cultured in complete assay medium for 72 h, then intracellular BCAAs were quantified using the BCAA-Glo Assay kit (Promega). Luminescence intensity was measured according to the manufacturer’s instructions, and results were normalized to cell count.

### Cut and Run followed by qPCR

Chromatin immunoprecipitation followed by RT-qPCR for the GLUT1 promoter region was performed using the CUT&RUN assay kit (CST #86652) following the manufacturer’s protocol. 100,000 MCF7 control or SLC7A5 cells with and without BCAT2 KD were used per reaction. The primary antibodies used for immunoprecipitation were 5 μg of rabbit mAb IgG isotype as a negative control (CST #66362) and 1 μg of rabbit Acetyl-Histone H3 (Lys27) (D534, 8173). Fifty picograms of spike-in control DNA (provided in kit) were added to each sample for normalization. DNA purification was performed by phenol/chloroform extraction followed by ethanol precipitation. RT-qPCR was performed using primers targeting a previously identified region of the GLUT1 promoter responsible for transcriptional activation^50,51^.

### Survival analysis of GSE186901 palbociclib cohort

Gene expression and clinical outcome data for patients with HR+/HER2− metastatic breast cancer treated with palbociclib monotherapy were obtained from NCBI Gene Expression Omnibus (GEO; accession GSE186901^34^). This cohort comprises 64 pre-treatment (baseline) tumor biopsy samples with paired RNA sequencing data and prospectively collected progression-free survival (PFS) endpoints. Expression values were provided as transcripts per million (TPM) and log₂-transformed [log₂(TPM+1)] prior to analysis. Only baseline biopsies were used; paired on-treatment samples were excluded.

For Kaplan-Meier survival analysis, patients were separated by median expression of SLC7A5. Survival curves were estimated using the Kaplan-Meier method and compared between groups using the two-sided log-rank test. Hazard ratios (HR) and 95% confidence intervals were estimated by Cox proportional hazards regression using the coxph function in the R survival package (v3.8-6).

For the SLC7A5+SLC2A1 composite analysis, a composite score was computed as the mean of log₂(TPM+1) values for SLC7A5 and SLC2A1, and entered as a continuous predictor in Cox regression. Spearman rank correlation was used to quantify the co-expression relationship between the two genes. The scatter plot visualization colors patients by above/below-median status in both genes simultaneously to illustrate the biological subgroups. All analyses were performed in R (v4.6.0). Survival curves were plotted using the survminer package (v0.5.2).

### Statistical analysis

Statistical analyses were conducted using Prism 10.0 (GraphPad Software) as previously described^52^. Briefly, statistical significances are shown as NS (not significant; P>0.05), *<0.05, **<0.01, ***<0.001, ****<0.0001. The sample size used in each experiment was not predetermined or formally justified for statistical power. To assess the statistical significance of a difference between two treatments, we used two-tailed Student’s t-tests. To assess the statistical significance of differences between more than two treatments, we used one-way or two-way ANOVA with Tukey’s multiple comparison test or Bonferroni’s multiple comparison test.

## Acknowledgements

This research was supported [in part] by the Intramural Research Program of the National Institutes of Health (NIH). The contributions of the NIH author(s) are considered Works of the United States Government. The findings and conclusions presented in this paper are those of the author(s) and do not necessarily reflect the views of the NIH or the U.S. Department of Health and Human Services. SKM received support from the Ludwig Center at Harvard University until 2022 and the Intramural Research Program of NIH (Grant: ZIABC012132). MUJO was supported by the NCI F99/K00 Pre-Postdoctoral Transition Fellowship and Black in Cancer/Emerald Foundation Career Transition Award. He is currently supported by the Dudley Rising Star Endowment at the University of Colorado. KN was supported by the NCI R35 Award and Breast Cancer Research Foundation (Joan Brugge PI). NK was supported by the Finnish Cultural Foundation (Suomen Kultturirahasto) Postdoc pool fellowship. TM is supported by the Ludwig Center at Harvard University.

## Author contributions

M.U.J.O. and S.K.M. designed, performed, and interpreted experiments and co-wrote the paper. J.C. and J.L. performed in vivo studies. M.H., K.K., and S.J. performed and contributed to metabolomics analyses. H.Y. performed RNA-sequencing analyses. K.N. contributed to cell line-based assays. N.K. and T.M. contributed to the design of drug treatment experiments.

---

**Extended Data Figure 1.**
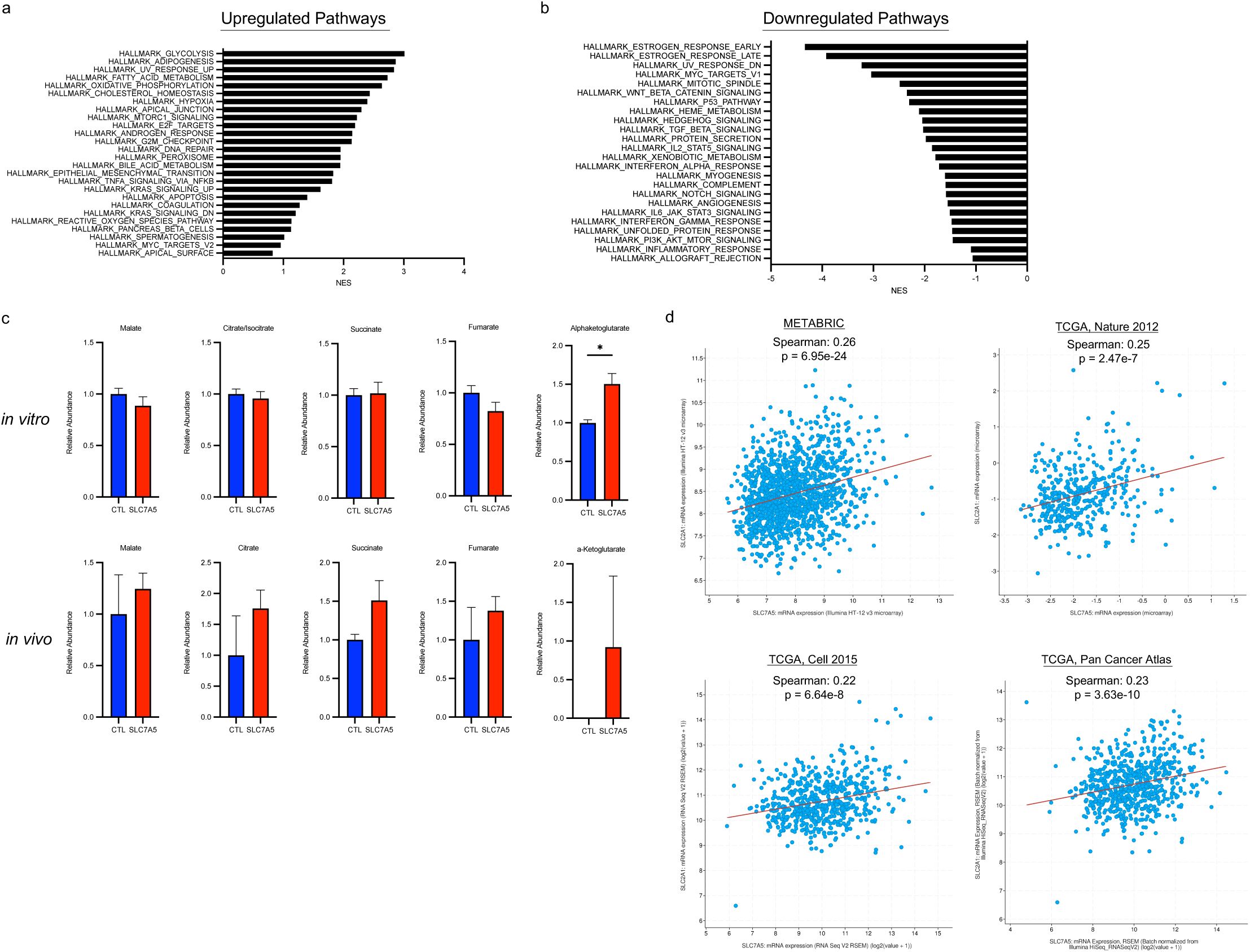
Normalized Enrichment Score (NES) bar plots for **a)** upregulated pathways and **b)** downregulated pathways in MCF7 SLC7A5 vs control cells. **c)** Steady state levels of TCA cycle metabolites from control and SLC7A5 overexpressing MCF7 cells grown in vitro and in vivo. **d)** Linear regression analysis demonstrating that SLC7A5 and GLUT1 mRNA significantly correlate in ER+ breast cancer patient samples (obtained from cBioportal).

**Extended Data Figure 2.**
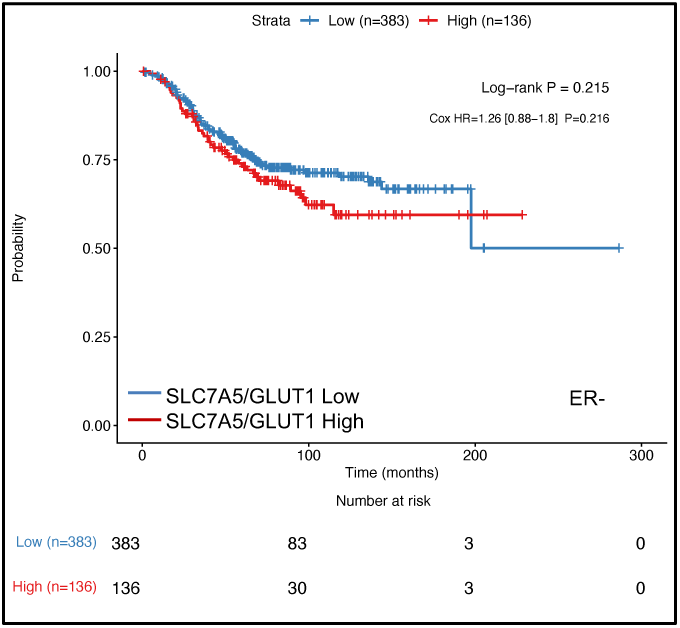
Overall survival in ER-tumors with high or low co-expression of SLC7A5 and GLUT1.

**Extended Data Figure 3.**
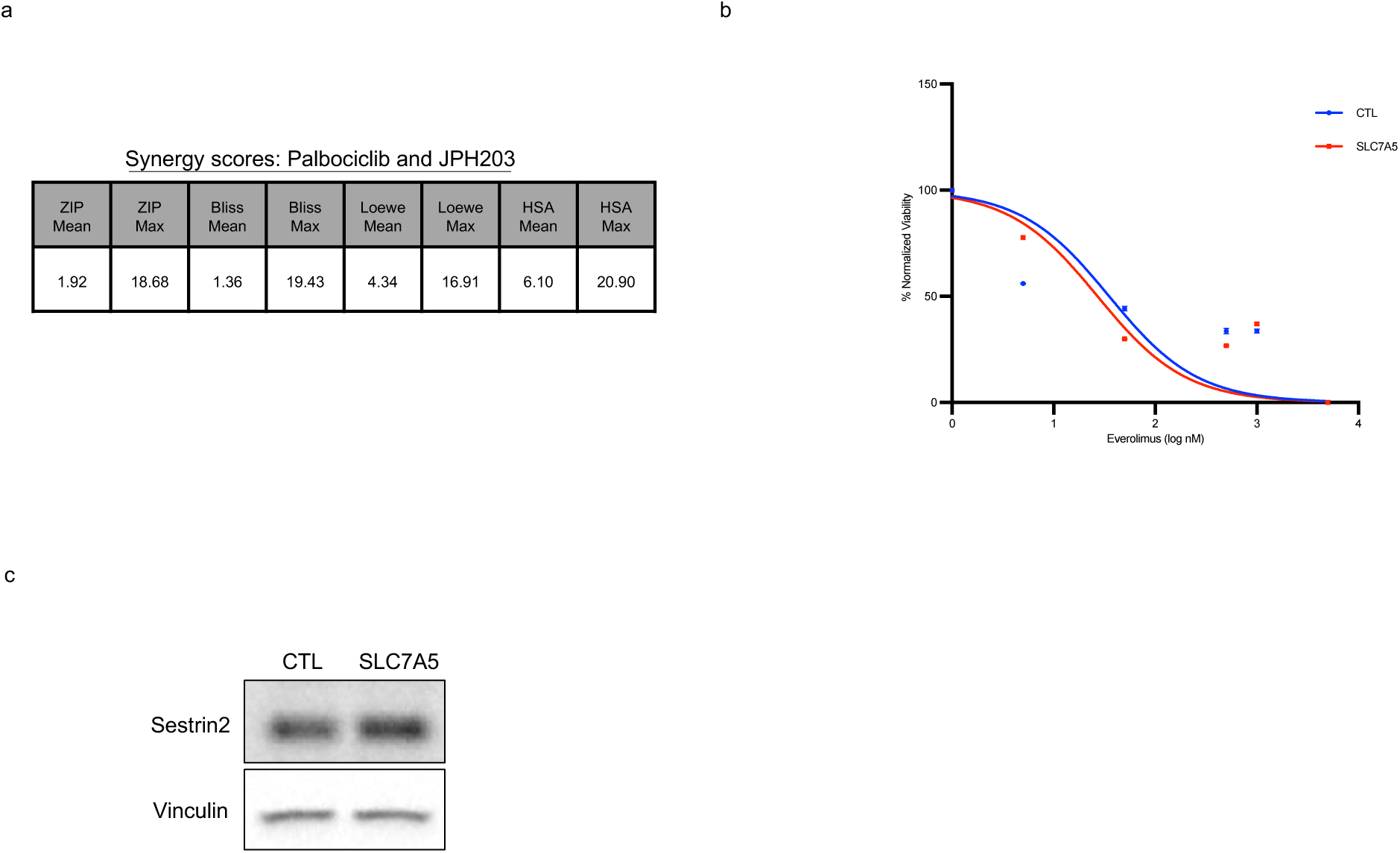
**a)** Synergy scores for combination treatment with palbociclib and JPH203 in MCF7 PalbR cells. Mean and max for each synergy model are shown. Analyses done using synergyfinder.org. **b)** Everolimus treatment in control and SLC7A5 overexpressing MCF7 cells. **c)** Immunoblot analysis of SESTRIN2 in control and SLC7A5 overexpressing MCF7 cells.

